# Impaired microglial phagocytosis promotes seizure development

**DOI:** 10.1101/2023.12.31.573794

**Authors:** Dale B. Bosco, Vaclav Kremen, Koichiro Haruwaka, Shunyi Zhao, Lingxiao Wang, Blake A. Ebner, Jiaying Zheng, Aastha Dheer, Jadyn F. Perry, Manling Xie, Aivi T. Nguyen, Gregory A. Worrell, Long-Jun Wu

**Author notes:** Correspondence, Department of Neurology, Mayo Clinic, 200 First Street SW, Rochester, MN 55905.

## Abstract

In the central nervous system, triggering receptor expressed on myeloid cells 2 (TREM2) is exclusively expressed by microglia and is critical for microglial proliferation, migration, and phagocytosis. TREM2 plays an important role in neurodegenerative diseases, such as Alzheimer’s disease and amyotrophic lateral sclerosis. However, little is known about the role TREM2 plays in epileptogenesis. To investigate this, we utilized TREM2 knockout (KO) mice within the murine intra-amygdala kainic acid seizure model. Electroencephalographic analysis, immunocytochemistry, and RNA sequencing revealed that TREM2 deficiency significantly promoted seizure-induced pathology. We found that TREM2 KO increased both acute *status epilepticus* and spontaneous recurrent seizures characteristic of chronic focal epilepsy. Mechanistically, phagocytic clearance of damaged neurons by microglia was impaired in TREM2 KO mice and the reduced phagocytic capacity correlated with increased spontaneous seizures. Analysis of human tissue from patients who underwent surgical resection for drug resistant temporal lobe epilepsy also showed a negative correlation between microglial phagocytic activity and focal to bilateral tonic-clonic generalized seizure history. These results indicate that microglial TREM2 and phagocytic activity may be important to epileptogenesis and the progression of focal temporal lobe epilepsy.

**One Sentence Summary:** Phagocytic activity of microglia may impact generalized seizure development within both mice and humans.

## INTRODUCTION

As the resident immune cells of the central nervous system (CNS), microglia have a diverse set of physiological functions and pathological roles (*1, 2*). Epilepsy is one of the most common neurologic diseases, affecting over 50 million people worldwide. Yet, the specific contribution of microglia to the development of seizures and focal epilepsy is less understood (*3*). What is known is that significant microglial activation occurs following chemotoxin seizure induction and that microglial activity is possibly beneficial during the acute phase of experimentally induced seizures (*3–5*). However, others have shown that disruption of native microglial function in and of itself can be epileptogenic (*6*). Clearly, more investigation is needed to determine what specific microglial factors can modulate seizures and the development of epilepsy.

Triggering receptor expressed on myeloid cell 2 (TREM2) is a surface receptor that, within the CNS, is exclusively expressed by microglia and influences a myriad of microglial functions(*7–9*). TREM2 can recognize a variety of ligands, such as fatty acids, Apolipoprotein-E, amyloid beta, and TDP-43 (*10–13*). TREM2 variants have been found in genome-wide association studies to increase the risk of Alzheimer’s disease (AD) (*14, 15*). Additionally, TREM2 has been related to other forms of neurodegenerative disease, such as amyotrophic lateral sclerosis (ALS) (*16, 17*). However, comparatively little is known about how TREM2 impacts seizures and epilepsy. In this context, patients diagnosed with polycystic lipomembranous osteodysplasia with sclerosing leukoencephalopathy (PLOSL), a rare condition directly related to deficiencies in TREM2 or its adaptor protein, DAP12, have been shown to develop seizures (*9*). While the initiator for seizures in PLOSL patients is unknown, the loss of TREM2-related microglia function is a likely contributor.

Here, we utilized constitutive TREM2 knockout (KO) mice within the intra-amygdala kainic acid (KA) seizure model, which can induce both acute *status epilepticus* (SE) and chronic focal epilepsy with spontaneously recurrent seizures (SRS) (*18*). We found that TREM2 KO increased the number of focal to bilateral tonic-clonic (FBTC) seizures, previously called secondarily generalized convulsions, both acutely and chronically. Additionally, TREM2 KO reduced microglial proliferation and impaired their ability to phagocytose damaged hippocampal neurons.

We then investigated resected human temporal lobe tissue from patients with a history of drug resistant focal epilepsy and found a negative correlation between the expression of microglial phagocytosis marker, CD68, and patient history of FBTC seizures. Patients displaying FBTC seizures are associated with more clinically severe epilepsy and less likely to benefit from surgical resection (*19*). Therefore, modulating microglial phagocytosis function via TREM2 signaling may attenuate seizure development and epilepsy progression, and provide a novel avenue for therapeutic intervention.

## RESULTS

### TREM2 KO alters kainic acid-induced seizure profiles

To investigate the role of microglial TREM2 in the pathological effects of seizure, we utilized the intra-amygdala KA administration model. FBTC seizure events (Racine score 4+) were quantified in both TREM2 KO and control mice. SE was terminated 45 mins after presentation of the first FBTC event to reduce animal loss. We found that while the time to the first FBTC event was similar between TREM2 KO and control mice, approximately 28 mins (Fig. 1A), TREM2 KO mice displayed a higher incident of FBTC events during the SE period (Fig. 1B, C). We also monitored animal survival for 7 days post-KA but found no statistical difference (Fig. 1D).

**Fig. 1:**
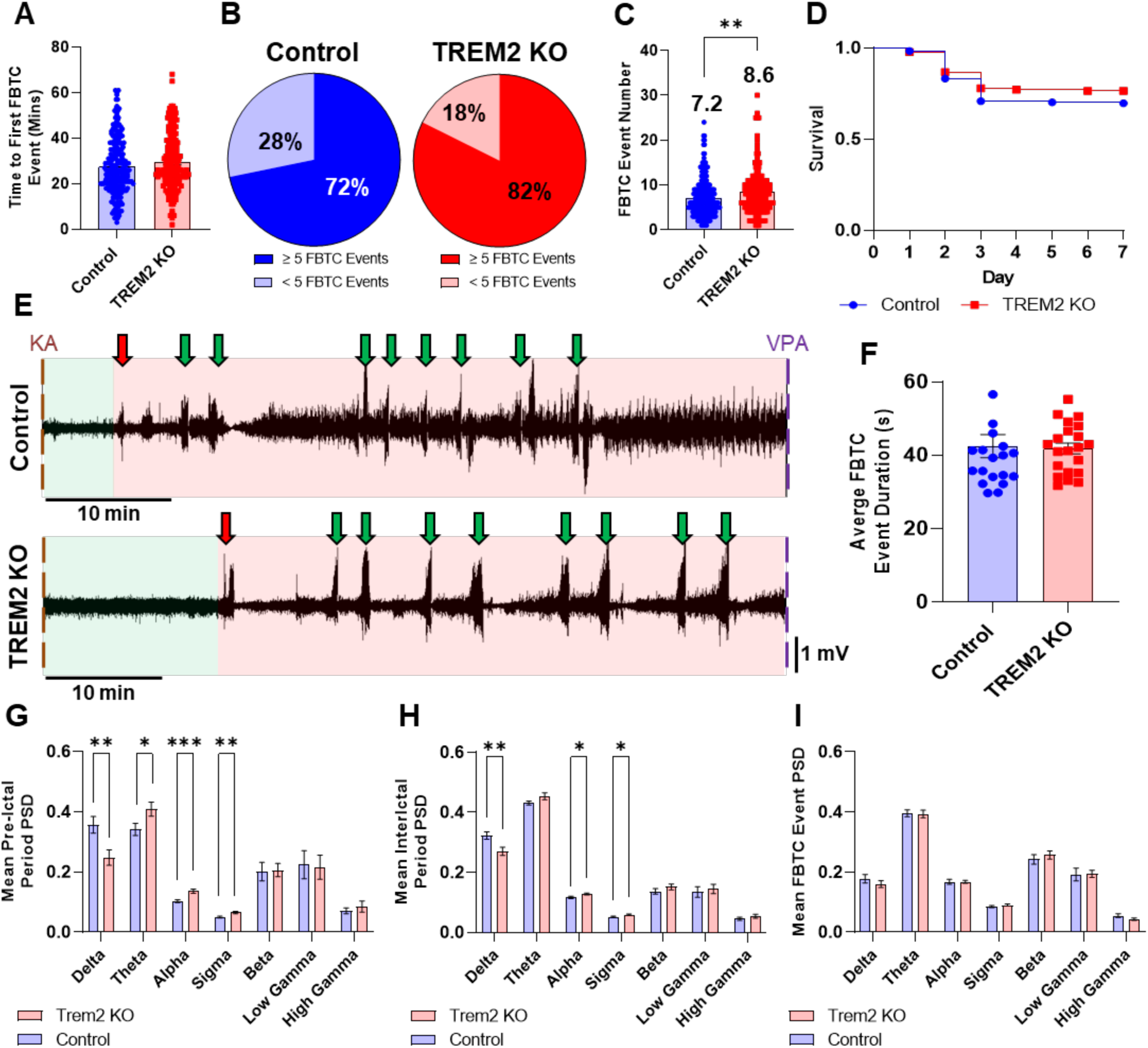
TREM2 alters EEG profiles of mice undergoing s*tatus epilepticus*. (A) No difference was observed between control and TREM2 KO in regards to when focal to bilateral tonic clonic (FBTC) events (Racine Score 4+) first occur. (B) TREM2 KO mice displayed a higher incidence of 5+ T-C events than controls. (C) The average number of T-C events was higher in TREM2 KO than controls. (D) TREM2 KO and control displayed similar post-KA survival. (E) Representative EEG profiles during s*tatus epilepticus.* Green area = pre-ictal period, Red area = ictal period, Red arrow indicates beginning of ictal period, Green arrows indicate FBTC events. (F) Average FBTC event duration was similar between TREM2 KO and controls. EEG profiles for each animal were then investigated for differences in individual frequency band power spectral density (PSD). Delta: 1-3 Hz; Theta: 3-9 Hz; Alpha: 8-12 Hz; Sigma: 12-15 Hz; Beta: 15-30 Hz; Low Gamma: 30-55 Hz; High Gamma: 65-110 Hz. (G) PSD evaluation for the pre-ictal period. (H) PSD evaluation for the interictal periods. (I) PSD evaluation for the aggregated FBTC events. All data presented as Mean ± S.E.M. N ≤ 18. Unpaired T-test with Welch correction was used for each comparison. * P < 0.05, ** P < 0.01, *** P < 0.001.

Given our initial observations, we wanted to further characterize the seizure profiles of control and TREM2 KO mice via cortical electroencephalographic (EEG) electrodes to analyze power spectral density (PSD) in conventional EEG bands (Fig. S1A). Overall, baseline electrographic characteristics were similar between groups, with minor changes in theta and high gamma bands (Fig. S1B, C). To determine how SE related EEG profiles were altered by TREM2 KO, we analyzed the EEG profiles following KA administration (Fig. 1E). Our analysis revealed that the average duration of all FBTC events was similar between TREM2 KO and control (Fig. 1F). Interestingly, when we compared PSD values between groups, we observed several differences. During the pre-ictal phase, the lack of TREM2 significantly altered the mean PSD of lower frequency bands (Fig. 1G). Dissecting out the FBTC seizures during the ictal period, we observed that electrographic differences are mainly restricted to the interictal segments, while the mean PSD for the FBTC seizures were nearly identical between groups (Fig. 1H, I). In total, our results show that TREM2 deficiency significantly alters the acute seizure behavioral and EEG profile in response to KA.

### TREM2 KO reduces microglial proliferation and reactivity following seizure induction

Our behavioral and EEG observations led us to next evaluate the effect of TREM2 KO on microglial response to KA-induced seizures. We found no significant difference in microglial number between groups under naive (baseline) conditions. However, following KA administration, significant differences emerged 7 days after intra-amygdala injection of KA in the hippocampal CA3 region and 14 days in CA1 region (Fig. 2A, B, Fig. S2A-C). Because hippocampal damage in this epilepsy model typically centers in the ipsilateral CA3, we focused further analysis mainly within this region. Next, we utilized bromodeoxyuridine (BrdU) treatment to investigate microglial proliferation in responses to seizures (Fig. 2C, D). We found significantly more BrdU positive microglia within controls than TREM2 KO mice at day 7 (Fig. 2E). Thus, our results suggest that TREM2 KO reduces microglial proliferation and microglial number following KA-induced seizures.

**Fig. 2:**
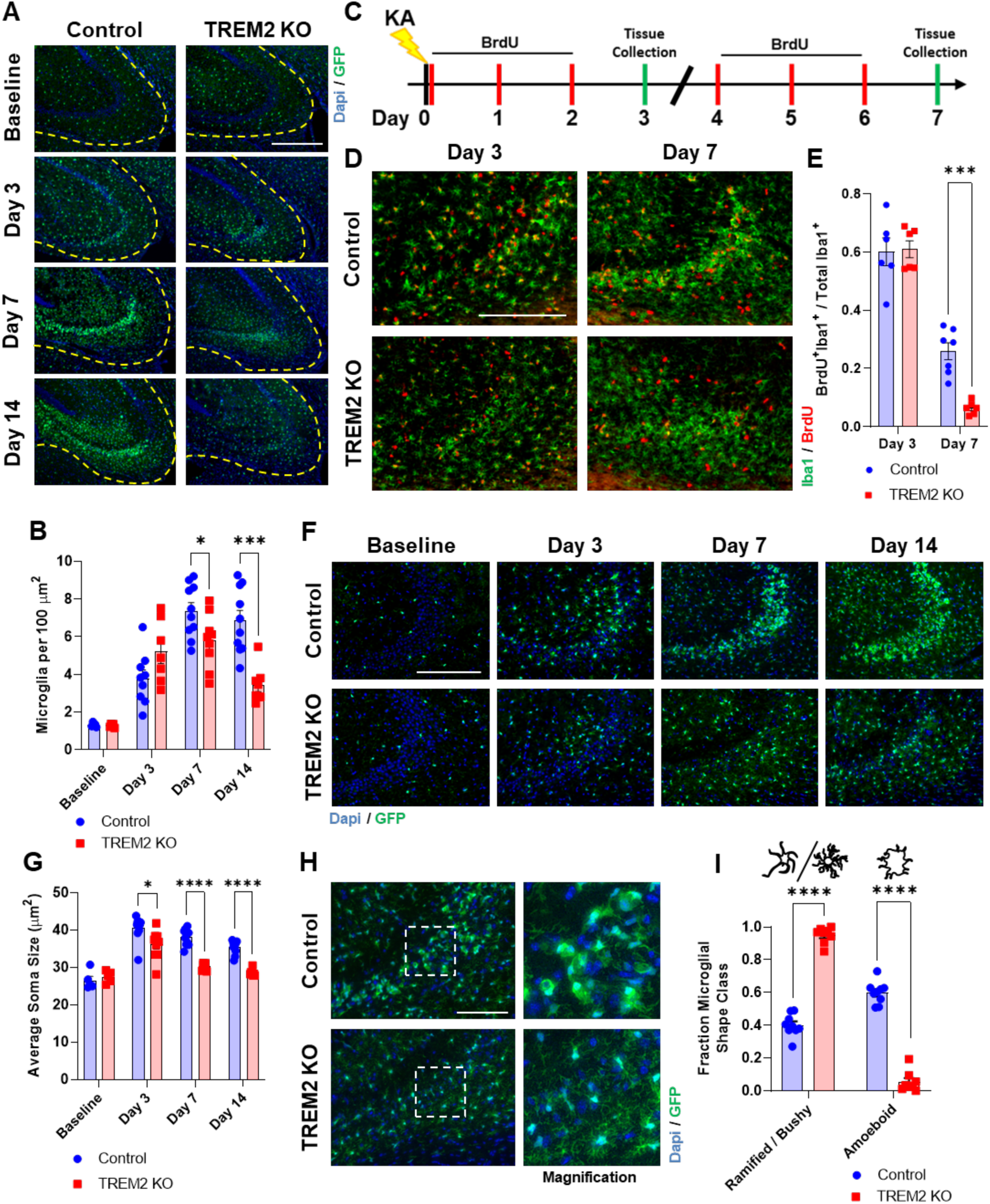
TREM2 affects microglial proliferation and morphology following IA-KA. (A) Representative images of the general CA3 region at baseline and 3, 7, 14 days post-KA administration. Scale bar = 400 µM. (B) Quantification of microglia density within the general CA3 region (Within the yellow dashed line). (C) Experimental timeline of BrdU administration and sample collection. Animals were administered BrdU twice daily on days 0-2, or days 4-6 post IA-KA. (D) Representative images of BrdU staining. Scale bar = 200 µm. (E) Quantification of BrdU^+^:IBA1^+^ cells divided by total IBA1^+^ cells. (F) Representative images of the CA3 region at baseline and 3, 7, 14 days post-KA administration. Scale = 200 bar µm. (G) Quantification of microglial soma size within the image field. (H) Representative images from tissues collected 7 days post-KA. Scale bar = 100 µm. (I) Quantification of the number of microglia displaying either ramified/bushy (small soma, long branched processes) or amoeboid (large soma, short processes) morphology in both control and TREM2 KO tissues. All data presented as Mean ± S.E.M. N = 5- 10 animals for each group. The quantification for 2-3 non-consecutive tissue sections were averaged for each N. Unpaired T-test with Welch correction was used for each comparison at each time point. * P > 0.05, *** P > 0.001, **** P > 0.0001.

Seizures also alter microglial morphology, where they transition from a ramified state to one that is more amoeboid with increased soma size (*20*). Interestingly, we found that while KA significantly increased the soma size of CA3 region microglia in controls, soma sizes remained close to basal levels in TREM2 KO (Fig. 2F, G). No consistent difference was observed in the CA1 region (Fig. S2D-F). Moreover, when looking at the CA3 region 7 days post-KA, nearly 60% of microglia in controls exhibited amoeboid morphology, while very few microglia in TREM2 KO mice adopted the amoeboid state typically observed with reactive microglia in various conditions (Fig. 2H, I). Together, these results indicate that TREM2 KO suppresses microglial morphology changes following KA-induced seizures, despite the higher number of acute FBTC seizures.

### TREM2 KO attenuates the clearance of apoptotic neurons

We next investigated how TREM2 KO affected neuronal survival following KA-induced seizure onset. NeuN staining revealed no appreciable difference at baseline or 3 days post-KA between groups. However, at 7 and 14 days post-KA, we observed that the loss of NeuN signal in the CA3 region was significantly greater in control than TREM2 KO (Fig. 3A, B). Co-localization of GFP with NeuN was also higher in control 7 days post-KA (Fig. 3C). In addition, at 7 days post-KA, microglia seem to be enveloping neurons in controls, while this was not observed in TREM2 KO mice (Fig. 3D). We observed minimal NeuN signal loss in the CA1 region in both groups across all timepoints (Fig. S3). To investigate the function of TREM2 in microglial phagocytosis after seizure, we first stained for lysosomal marker CD68. Indeed, there is a significant decrease in CD68 expression at later timepoints in TREM2 KO in the CA3 region (Fig. 3E, F). The effect upon phagocytosis was validated *in vitro*, where TREM2 KO microglia displayed less capacity for pHrodo red labeled apoptotic N2a cells (Fig. 3G-I).

**Fig. 3:**
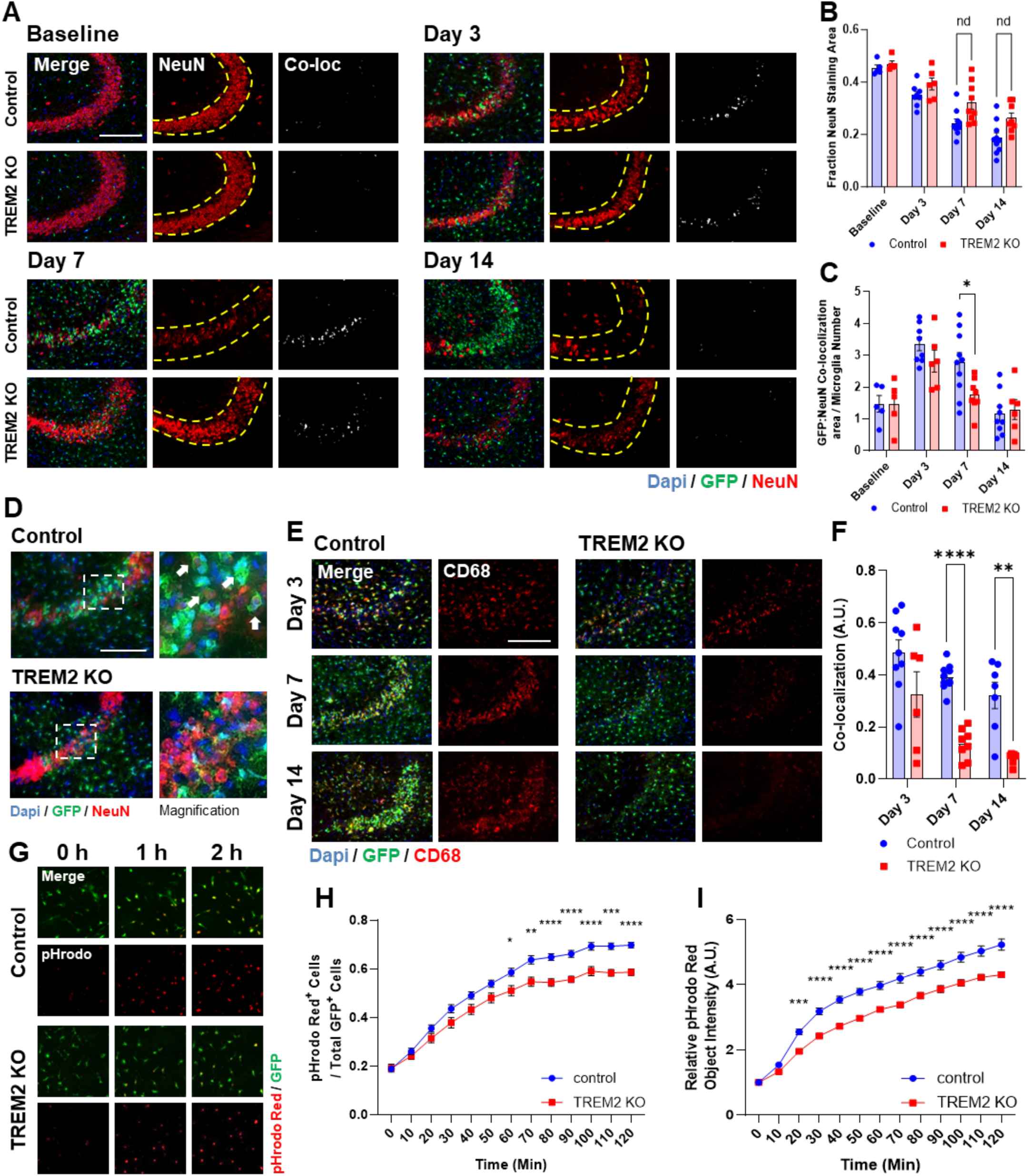
TREM2 KO affects neuronal atrophy following IA-KA administration. (A) Representative CA3 region images of NeuN staining and GFP:NeuN co-localization. Scale bar = 200 µm. (B) Positively stained NeuN area divided by total area (between yellow dashed lines). (C) GFP:NeuN co-localization divided by microglia number within the image field. (D) Representative images from 7 days post-KA. Scale bar = 100 µm. There are indications of microglia engulfing neurons following IA-KA in controls, but not TREM2 KO. Magnification, White arrows. (E) Representative images of CD68 staining within the CA3 region. Scale bar = 200 µm. (F) GFP and CD68 staining signal co-localization. (G) Representative images of primary microglia incubated with apoptotic N2a cells over time. Scale bar = 200 µm. (H) pHrodo red positive microglia relative to total microglia number. (I) Average pHrodo red object integrated intensity normalized to time 0 h. All data presented as Mean ± S.E.M. For B, C, F N = 5-10 for each group. Images from 2-3 non-consecutive tissue sections were averaged for each N. Unpaired T-test with Welch correction was used for each comparison. For H, I N = 11-12 replicates from 2 independent experiments. Four image fields averaged for each N. 2-way ANOVA followed by post-hoc multiple comparisons were used for each comparison. * P < 0.05, ** P < 0.01, *** P < 0.001, **** P < 0.0001.

We next investigated if the reduction in phagocytic capacity was related to recognition of apoptotic markers. Phosphatidylserine (PS) exposure acts as an “eat-me” signal when a cell undergoes apoptosis (*21*). It has been suggested that TREM2 can sense apoptotic cells via PS recognition (*11, 12*). To investigate this, we treated mice with PSVue 550, a PS specific dye, to determine PS exposure on the outer membrane after KA. We found that there was no significant difference in PSVue^+^ cell number, at 3 or 7 days post-KA (Fig. 4A-B). This is consistent with Fluoro-Jade C (FJC) staining which specifically labels dead or dying neurons (Fig. S3C-D). However, there was substantially less co-localization with GFP in TREM2 KO than control (Fig. 4C). We further investigated the amount of contact that was occurring between microglia and individual PSVue^+^ cells (Fig. 4D). We categorized contacts into three groups based on the overlap of GFP signal around the outer edge of PSVue^+^ cells, little to no contact (<20%), contact (20 – 90%), and engulfment (>90%) (Fig. 4E). Consistent with our co-localization results, we found that at 3 days post-KA, the percentage of engulfed PSVue^+^ cells was similar between control and TREM2 KO (3.6% vs 4.6%, respectively) (Fig. 4F, G). However, at 7 days post-KA, engulfment accounts for 25% of all contacts in controls, but only 5.2% in TREM2 KO (Fig. 4H, I). These results indicate that loss of TREM2 function interferes with apoptotic cell recognition and phagocytosis.

**Fig. 4:**
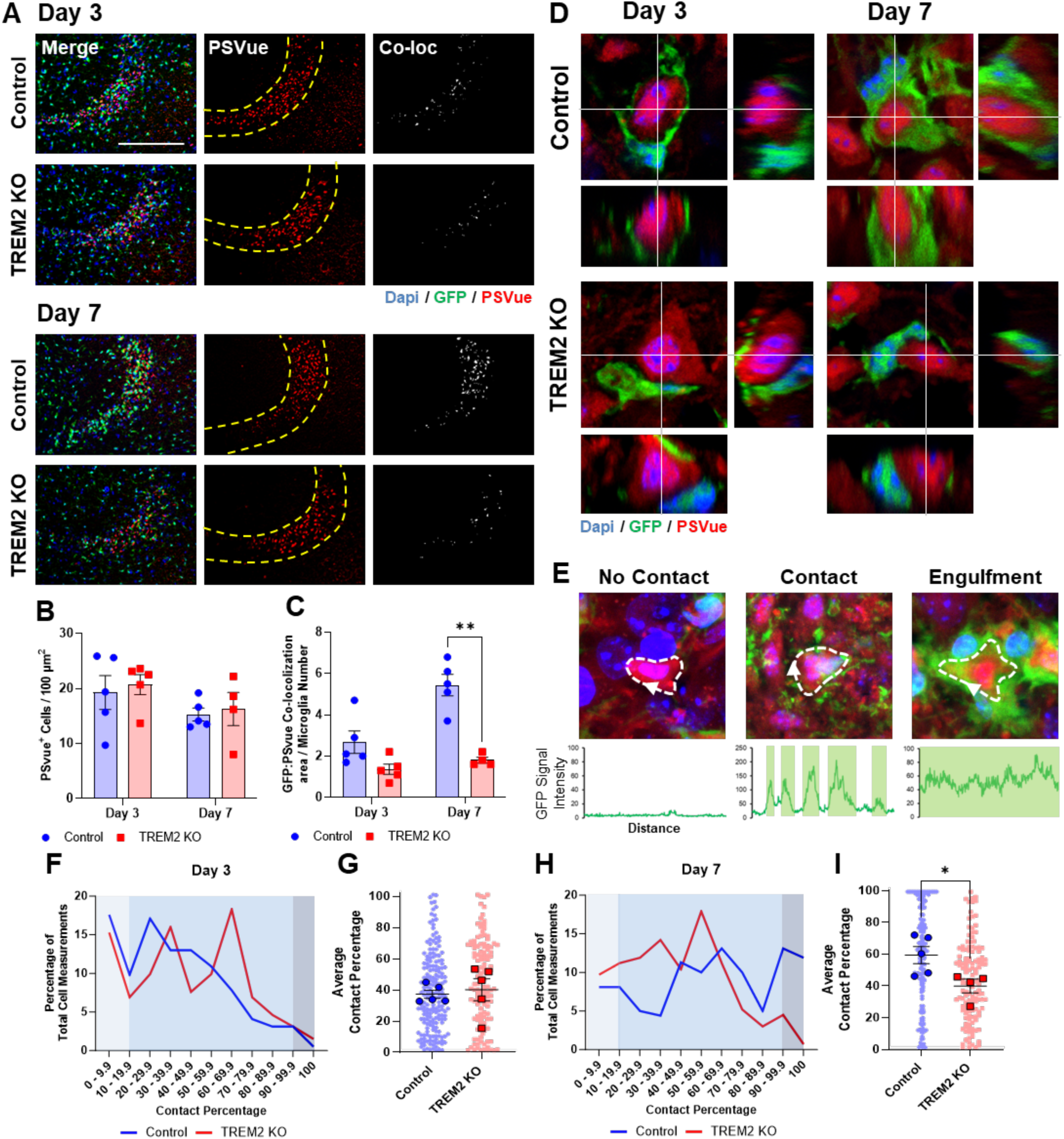
TREM2 KO affects microglial response to apoptotic neurons. (A) Representative images of PSVue staining and GFP:PSVue co-localization within the CA3 region. Scale bar = 200 µm. (B) PSVue^+^ cells divided by total area (between yellow dashed lines). (C) GFP:PSVue co-localization area divided by microglia number. (D) Representative Z direction images of microglia interacting with PSVue^+^ cells. Scale bar = 6 µm. (E) Representative images and intensity plots for various levels of microglial interaction along the outer edge of PSVue^+^ cells. (F) Binned 3 days post-KA cellular contact measurements. (G) Average contact percentage at 3 days post-KA. (H) Binned 7 days post-KA cellular contact measurements. (I) Average contact percentage at 7 days post-KA. All data presented as Mean ± S.E.M. For B and C, N = 4-5 for each group. Images from 2-3 non-consecutive tissue sections were averaged for each N. For G, I contact measurements from 10-50 cells per animal were averaged for each N. Unpaired T-test with Welch correction was used for all comparisons. * P < 0.05, ** P < 0.01.

### RNAseq analysis reveals key differences in gene expression profiles

Delving deeper into the pathological differences introduced by TREM2 KO, we performed RNAseq analysis on microglia isolated from the general region of KA injection (Table S1). In addition to the baseline condition, we investigated 7 days post-KA, as this is when we observed the most substantial difference between groups. We found a significant increase in differentially expressed genes (DEGs) following KA administration (Fig S4, Tables S2-5). A volcano plot highlights differences in expression between control and TREM2 KO with a Log_2_ fold change ≥1 (Fig. 5A). At this threshold, approximately 300 genes were significantly upregulated, while about 200 genes were downregulated in TREM2 KO mice when compared with control. Network analysis of these 2 gene groups revealed that a large proportion of DEGs were connected. This suggests that loss of TREM2 function interferes with highly interdependent pathways (Fig. 5B). Gene ontology (GO) analysis revealed these connections were related primarily to cellular adhesion and cell cycle biological processes (Fig. 5C). Of the cell cycle-related genes identified, 100% were up-regulated over baseline in controls, but only 17% were up-regulated over baseline in TREM2 KO and to not the same degree (Fig. 5D). These observations are consistent with our microglia proliferation results showing reduced microglia proliferation and number in TREM2 KO mice when compared with control at 7 days after KA injection.

**Fig. 5:**
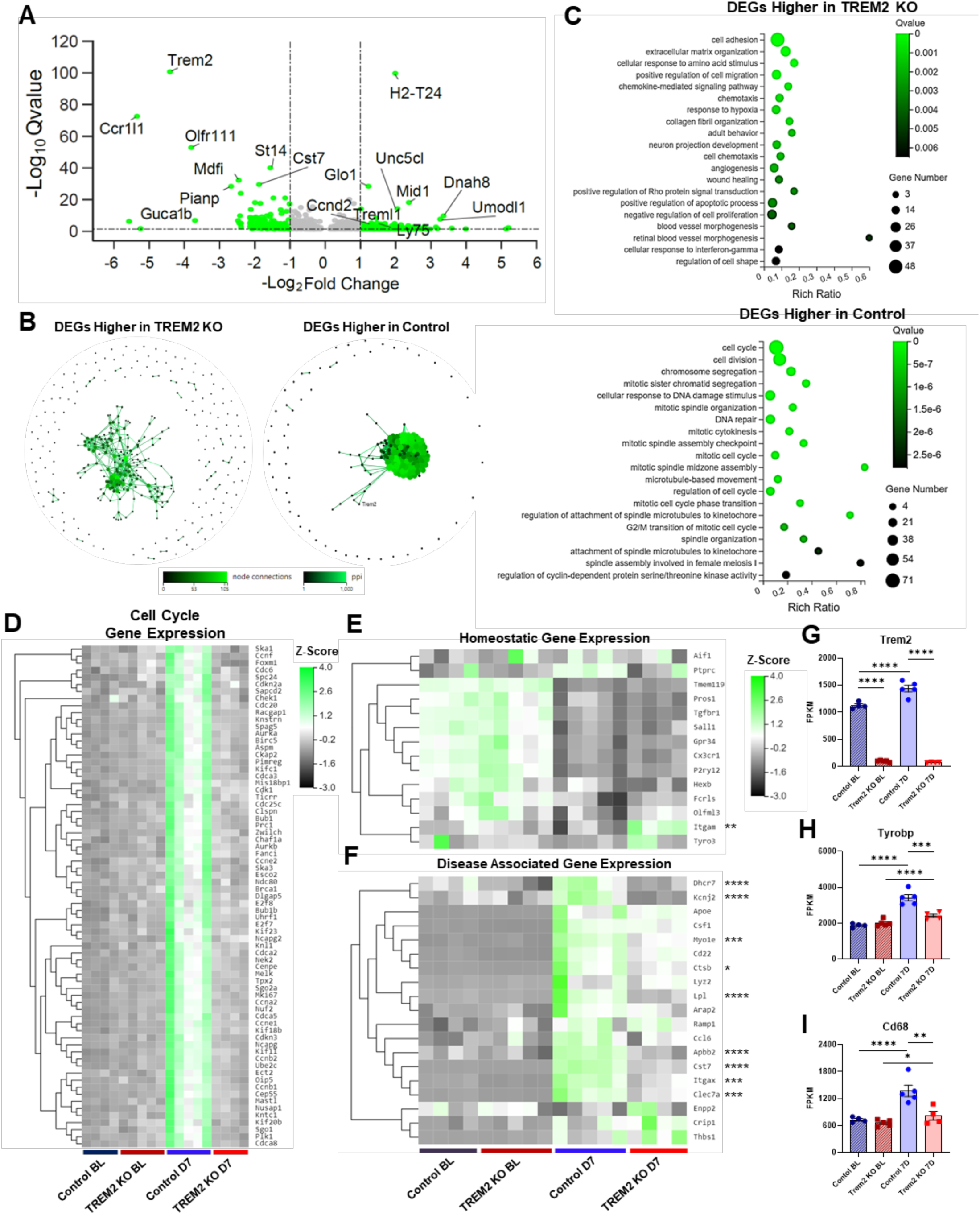
Comparison of TREM2 KO and control gene expression profiles. (A) Volcano plot illustrating differentially expressed genes (DEGs) with a log_2_ ≥ 1. (B) Network analysis of identified DEGs with expression log_2_ ≥ 1 higher in TREM2 KO or control. (C) Biological process gene ontology (GO) of identified DEGs. (D) Heat map of identified cell cycle related DEGs at both baseline and 7 days post-KA. (E) Heat map of homeostatic microglial genes. (F) Heat map of disease associated microglial genes. Comparisons in gene expression for (G) Trem2, (H) Tyrobp (DAP12) and (I) Cd68. FPKM = Fragments per kilobase of transcript per million fragments mapped. Analysis and comparison between gene sets was carried out via DESeq2. For E, F, asterisks indicate statistically significate differences in expression between TREM2 KO and control at 7 days post-KA. For all comparisons significance is expressed by Q value. * Q < 0.05, ** Q < 0.01, *** Q < 0.001, **** Q < 0.0001.

We have shown previously that following seizure induction, microglia alter homeostatic gene expression (*22*). Interestingly, we found that TREM2 KO did not significantly disrupt expression changes to classically termed homeostatic genes when compared to control (Fig. 5E). We next wanted to determine if TREM2 KO affected the expression of disease-associated microglia (DAM) genes. It is known within AD literature that TREM2 can influence the adoption of the DAM phenotype (*23, 24*). We found that while TREM2 KO may not significantly affect homeostatic gene regulation following KA administration, it does impact genes associated with microglial reactivity (9/19 genes) (Fig. 5F). Thus, TREM2 KO microglia may be locked in a phenotype that is between homeostatic microglia and reactive microglia, compromising their ability to perform necessary functions. We also found that there was an approximate 30% increase in TREM2 expression in controls following KA administration (Fig. 5G). We also observed that increases in expression of the TREM2 binding partner Tyrobp (DAP12) were also higher in control (Fig. 5H). Lastly, we found that while CD68 expression is significantly increased over baseline in control, TREM2 KO displayed only a minimal change in expression (Fig. 5I). This again is consistent with our previously outlined results on TREM2 function in microglial phagocytosis after seizures.

### TREM2 KO affects epileptogenesis and spontaneous recurring seizure development

Intra-amygdala KA administration not only induces acute SE response, but also spontaneous recurrent seizures (SRS) within a rather short timeframe (*18*). Our results so far have demonstrated that TREM2 deficiency impaired microglial phagocytosis and neuronal clearance. The next question is what the consequence of this defect is for epileptogenesis. We found that while both genotypes exhibit similar SRS events, TREM2 KO mice trended towards a higher incidence of SRS presentation during the five-day recording period (day 7-11 post-KA) (Fig. 6A, B). Moreover, of the mice that displayed SRS, TREM2 KO had a higher frequency of events than control (Fig. 6C, D). SRS duration and time (day/night cycle) of presentation was not significantly affected (Fig. 6E, F). Finally, we investigated interictal PSDs at 7 days post-KA, when we observed the greatest difference between groups. We found a similar pattern of differences as the acute SE pre-ictal and interictal EEG profiles (Fig. 6G). This indicates that seizure-induced EEG profile differences are maintained for days after KA administration. Moreover, when normalizing to genotypic baselines, it seems that these differences were predominantly related to the EEG patterns of the TREM2 KO mice (Table 1). Consequently, loss of TREM2 function not only affects the response of microglia after KA but also epileptogenesis.

**Fig. 6:**
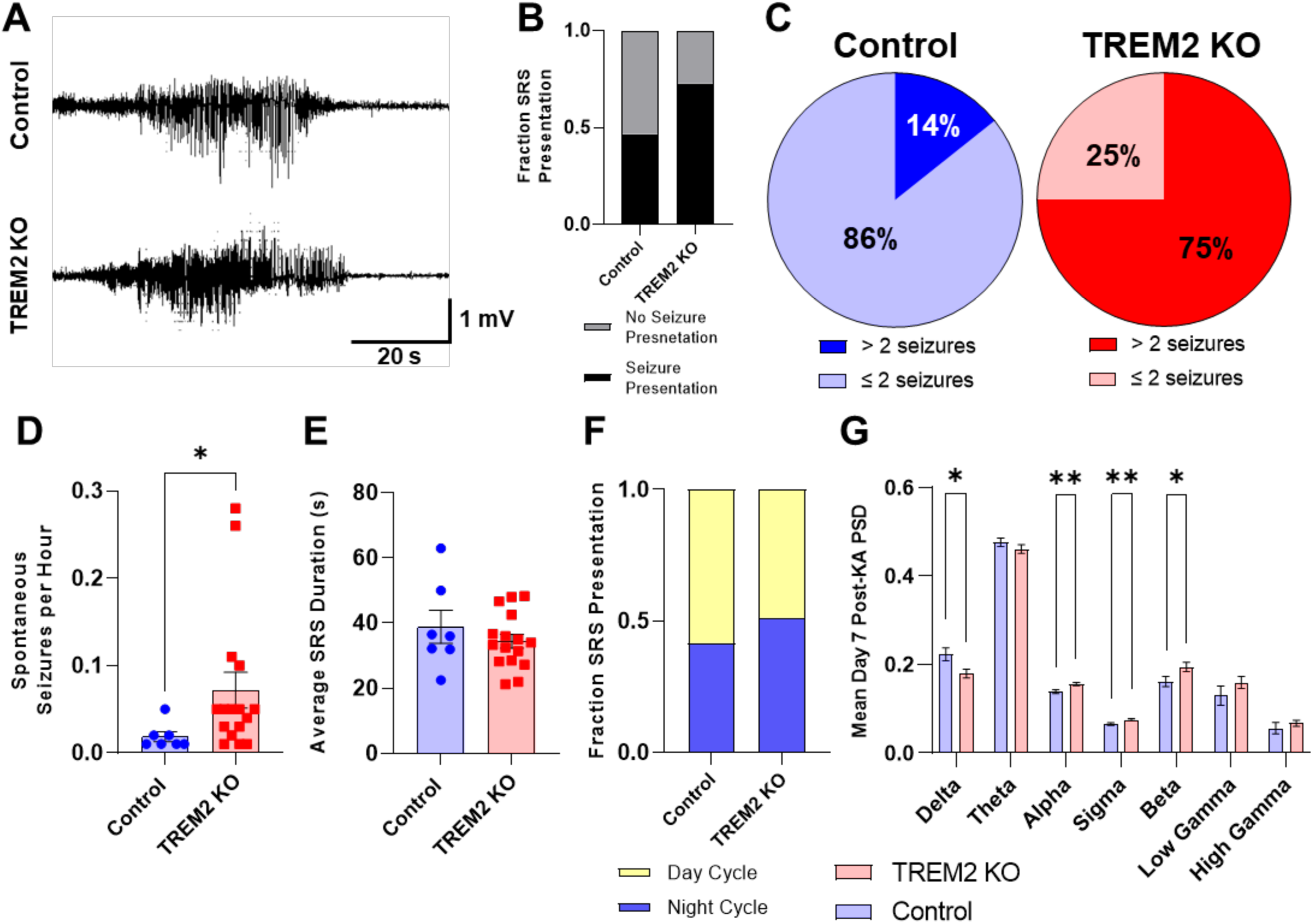
TREM2 KO increases spontaneous recurrent seizure frequency. (A) Representative EEG traces captured during spontaneous recurrent seizure (SRS) events. (B) Fraction of animals that presented SRS versus the total number of animals enrolled. (C) Percentage of animals presenting SRS activity displaying +2 SRS events during the recording period (Day 7-11 post-KA). (D) SRS frequency. (E) Average SRS event duration. (F) No difference was observed as to when SRS events occurred during the housing day/night cycle. (G) PSD evaluations of 3 consecutive hours of non-FBTC event containing EEG recordings collected 7 days post-KA. All data presented as Mean ± S.E.M. N ≤ 7 for each group. Unpaired T-test with Welch correction was used for each comparison. * P < 0.05, ** P < 0.01.

**Table 1:** Relative EEG band values for various periods.

| **Control** |  |  |  |  |  |  |
| --- | --- | --- | --- | --- | --- | --- |
| **Mean ± S.E.M.** | **Baseline** | **Pre-Ictal** | **Ictal** | **Interictal** | **T-C Events** | **7D Post-KA** |
| **Delta** | 1.000 ± 0.099 | 1.704 ± 0.131 | 1.445 ± 0.059 | 1.543 ± 0.059 | 0.849 ± 0.068 | 1.065 ± 0.070 |
| **Theta** | 1.000 ± 0.014 | 0.712 ± 0.417 | 0.905 ± 0.017 | 0.899 ± 0.013 | 0.825 ± 0.024 | 0.994 ± 0.019 |
| **Alpha** | 1.000 ± 0.051 | 0.704 ± 0.034 | 0.828 ± 0.023 | 0.796 ± 0.028 | 1.143 ± 0.057 | 0.949 ± 0.032 |
| **Sigma** | 1.000 ± 0.075 | 0.746 ± 0.047 | 0.836 ± 0.026 | 0.783 ± 0.032 | 1.278 ± 0.059 | 0.985 ± 0.043 |
| **Beta** | 1.000 ± 0.082 | 1.205 ± 0.185 | 0.879 ± 0.047 | 0.816 ± 0.057 | 1.450 ± 0.094 | 0.968 ± 0.069 |
| **Low Gamma** | 1.000 ± 0.101 | 1.550 ± 0.325 | 0.925 ± 0.102 | 0.934 ± 0.122 | 1.328 ± 0.148 | 0.901 ± 0.148 |
| **High Gamma** | 1.000 ± 0.109 | 1.354 ± 0.186 | 0.862 ± 0.097 | 0.864 ± 0.106 | 1.047 ± 0.117 | 1.087 ± 0.243 |
| **TREM2 KO** |  |  |  |  |  |  |
|  | **Baseline** | **Pre-Ictal** | **Ictal** | **Interictal** | **T-C Events** | **7D Post-KA** |
| **Delta** | 1.000 ± 0.071 | 1.119 ± 0.118 | 1.214 ± 0.061 | 1.223 ± 0.062 | 0.722 ± 0.051 | 0.813 ± 0.046 |
| **Theta** | 1.000 ± 0.018 | 0.919 ± 0.053 | 0.993 ± 0.024 | 1.020 ± 0.026 | 0.884 ± 0.028 | 1.038 ± 0.023 |
| **Alpha** | 1.000 ± 0.034 | 0.966 ± 0.052 | 0.948 ± 0.027 | 0.909 ± 0.023 | 1.192 ± 0.035 | 1.113 ± 0.027 |
| **Sigma** | 1.000 ± 0.054 | 0.962 ± 0.061 | 0.920 ± 0.037 | 0.872 ± 0.034 | 1.332 ± 0.055 | 1.128 ± 0.037 |
| **Beta** | 1.000 ± 0.072 | 1.070 ± 0.115 | 0.819 ± 0.054 | 0.785 ± 0.052 | 1.336 ± 0.072 | 1.014 ± 0.054 |
| **Low Gamma** | 1.000 ± 0.079 | 1.151 ± 0.212 | 0.755 ± 0.075 | 0.772 ± 0.083 | 1.037 ± 0.062 | 0.853 ± 0.070 |
| **High Gamma** | 1.000 ± 0.093 | 1.173 ± 0.253 | 0.688 ± 0.082 | 0.744 ± 0.089 | 0.595 ± 0.055 | 0.938 ± 0.097 |
| **Ratio** |  |  |  |  |  |  |
|  | **Baseline** | **Pre-Ictal** | **Ictal** | **Interictal** | **T-C Events** | **7D Post-KA** |
| **Delta** | N/a | 0.656 | 0.840 | 0.792 | 0.850 | 0.764 |
| **Theta** | N/a | 1.290 | 1.097 | 1.135 | 1.072 | 1.043 |
| **Alpha** | N/a | 1.374 | 1.145 | 1.142 | 1.043 | 1.172 |
| **Sigma** | N/a | 1.290 | 1.100 | 1.113 | 1.042 | 1.145 |
| **Beta** | N/a | 0.888 | 0.932 | 0.962 | 0.921 | 1.048 |
| **Low Gamma** | N/a | 0.743 | 0.816 | 0.827 | 0.781 | 0.946 |
| **High Gamma** | N/a | 0.866 | 0.798 | 0.861 | 0.568 | 0.863 |

### Microglial phagocytosis within tissue resections from humans with drug resistant temporal lobe epilepsy

We finally wanted to determine if the observed effects of microglial phagocytic activity upon seizure presentation extended to human temporal lobe epilepsy (TLE). Temporal lobe and hippocampal tissue sections were obtained from patients that underwent unilateral anterior temporal lobe and amygdala-hippocampal resection for drug resistant TLE. Selected patients presented pathological markers of mesial temporal lobe epilepsy with no known history of brain cancer or focal cortical dysplasia (*25*). Sections from both areas were immunofluorescently stained for Iba1 and CD68, as indication for microglial phagocytosis (Fig. 7A). For hippocampal sections we found that the number of Iba1^+^ cells was highest near the dentate gyrus (DG), while only proximal white matter (WM) Iba1^+^ cells displayed any significant CD68 expression (Fig. 7B, C). Looking at temporal lobe sections, we observed substantially more Iba1^+^ cells and CD68 expression in WM regions than grey matter (GM) (Fig. 7D, E).

**Fig. 7:**
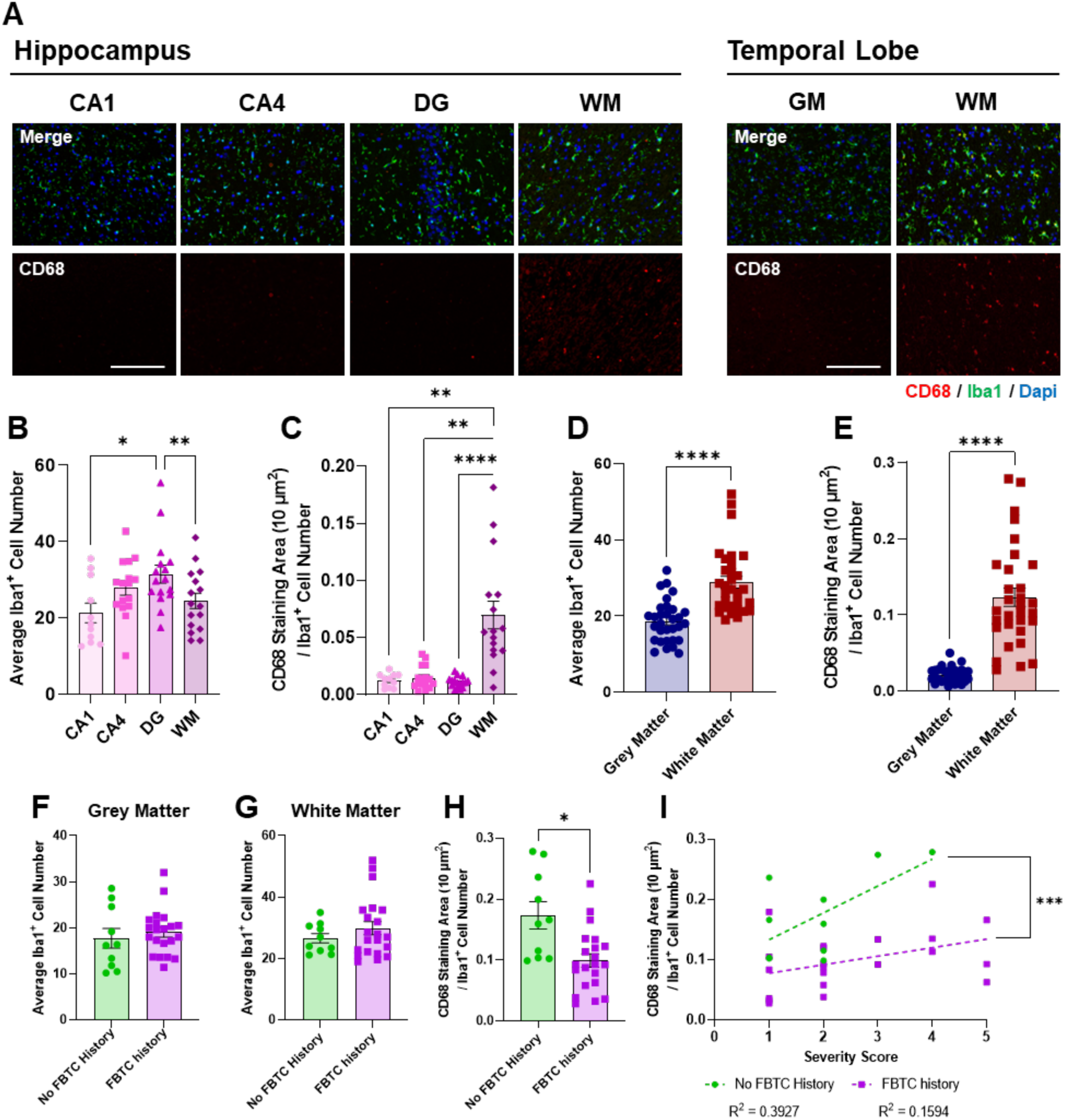
CD68 expression within human epilepsy derived tissues sections. (A) Representative images of hippocampal CA1, CA4, dentate gyrus (DG) and proximal white matter (WM) and temporal lobe grey matter (GM) and WM. Scale bar = 200 µm. (B) Analysis of Iba1^+^ cell number in the hippocampus. (C) CD68 staining area per Iba1^+^ cell in the hippocampus. (D) Iba1^+^ cell number in the temporal lobe. (E) CD68 staining area per Iba1^+^ cell in the temporal lobe. Sorting results by generalized seizure history, no difference was found in Iba1^+^ cell number in temporal lobe (F) GM and (G) WM. (H) Temporal lobe WM CD68 expression was found to be higher in patients with no known history of generalized seizures. (I) Plotting the sorted data in H against severity score (0 = 0, 1 = ≤1, 2 = 2-4, 3 = 5-7, 4 = 8-12, 5 = 13≤ monthly focal seizures) revealed that patients without generalized seizures had higher CD68 expression at every severity level. All data presented as Mean ± S.E.M, except for I. N ≤ 10 patients per group. 2-5 image fields per region averaged for each N. For B, C 1-way ANOVA was used for each comparison. For F-H Unpaired T-test with Welch correction was used for each comparison. For I difference is based on linear regression elevation. * P < 0.05, ** P < 0.01, *** P < 0.001, **** P < 0.0001.

We next determined whether a patient’s documented history of FBTC seizures related to CD68 expression by Iba1^+^ cells in the temporal lobe, as this is where we observed the highest expression. We found that while FBTC presentation did not affect GM or WM Iba1^+^ cell number, WM CD68 expression was, on average, higher in patients without a documented history of FBTC (Fig. 7F- H). We also organized patients based on the number of monthly focal seizures the patients presented (severity scores). There was no apparent correlation between severity score and disease duration or age at onset, nor did it matter if the tissue sections originated from the left or right temporal lobes, or male or female patients (Fig. S4). When we plotted WM CD68 expression against severity score, we found that at every severity score, patients without a documented FBTC history had higher average CD68 expression (Fig. 7I). This is in line with our results regarding TREM2 KO, where we also observed a negative relationship between apparent microglial phagocytic activity and FBCT events. Taken together, phagocytic activity by microglia may be protective in regard to the presentation of FBTC events in both mice and humans.

## DISCUSSION

While the development of seizures and epilepsy has been intensely investigated, there are many aspects of these conditions that remain unknown, specifically the roles glial cells play. Here, we sought to determine how microglial TREM2 contributes to the development of seizures and epilepsy. We observed that TREM2 KO reduces the reactivity and phagocytic capacity of microglia in response to seizures and neuronal damage. This is in turn correlated with exacerbated KA-induced SRS development. We also observed that CD68 expression, an indirect marker of phagocytic activity, negatively correlated with FBTC seizures in human TLE patients. Focal seizures that propagate more widely through the brain generating FTBC seizures are more dangerous for patients and known to be associated with sudden unexplained death in epilepsy (SUDEP) (*26*). Taken together our results indicate that modulation of microglial phagocytic capacity could impact generalized seizure burden.

### TREM2 modulates acute seizure development and EEG profiles

We found that TREM2 deficiency increased acute seizures and EEG profile in response to KA. Considering the important role of microglia in regulation of neuronal activity, it is possible that microglial TREM2 modulates the interaction of microglia with hyperactive neurons (*27*). This can be seen in both the pre-ictal and interictal periods during *status epilepticus* where lower frequency bands are altered in TREM2 KO mice compared to controls. It has also been suggested within the AD literature that TREM2 deficiency can affect epileptiform activity (*28, 29*). However, these studies did not investigate the effect upon generalized seizure presentation nor a specific role for microglia in the context of epilepsy. Other studies have demonstrated that microglial processes are attracted to hyperactive neurons and can possibly dampen seizure activity (*30, 31*). As such, TREM2 deficiency may reduce the threshold for neuronal hypersynchronization resulting in higher generalized seizure burden. This could also explain why depletion of microglia affects seizure burden, because again the proposed neuronal dampening activity would be impaired. Consistent with the idea of neuroprotective function, we and others have shown microglial depletion will increase seizure burden and severity (*5, 32, 33*). Microglia and TREM2 have also been implicated in synaptic pruning and maintaining synaptic function (*34–36*). Additionally, investigations into microglial DAP12, the principal binding partner of TREM2, showed that DAP12 deficiency will alter synaptic function and enhanced long-term potentiation (*37*). Considering that the animals utilized within this study are general knockouts, this effect may introduce some level of baseline developmental difference in regard to circuit formation. These observations may also relate to the development of seizures and epilepsy within patients suffering from PLOSL, a disorder directly related to the genetic loss of TREM2 function or its adaptor protein DAP12 (*38*). Regardless, further investigation is needed to determine the exact mechanism underlying how microglial TREM2 could participate in the regulation of neuronal circuits and the dampening of neuronal hyperactivity acutely.

### TREM2 affects microglial response to KA-induced pathology in a temporal fashion

During the course of our investigation, we found no significant differences in microglial number at baseline or 3 days post-KA. This contrasts with Zheng *et al.,* which found that at 3D post-KA proliferation was reduced in TREM2 KO mice (*39*). This difference may arise from the fact that their study utilized a different KA model. Nevertheless, when comparing microglia number at later timepoints, we found significantly fewer microglia in TREM2 KO than control. Both BrdU analysis and RNAseq results confirmed that proliferation is affected by TREM2 KO. Additionally, while down-regulation of homeostatic genes was not altered by TREM2 KO, up-regulation of diseased associated gene expression was mitigated. The differences in microglial response between early and later timepoints could be explained by the fact that while TREM2 is an important receptor for microglial function, it is not their only receptor. Multiple studies reported that microglial response to seizures and epilepsy can be modulated by targeting a plethora of receptors including P2Y12, CX3CR1, and CSF1R (*40–43*). Microglial CSF1R and CX3CR1 activation after seizure is a key component of their response. ATP is known to be released from hyperexcited neurons during seizures, which can activate microglia via P2Y12 and P2X7 (*30, 44*). Moreover, direct modulation of microglial mTOR signaling has been shown to be epileptogenic (*6*). Consequently, it is possible that while other receptors are responsible for initial responses by microglia following KA, sustained response and completion of their phenotypic transition is predicated upon TREM2 signaling.

From removal of invading bacteria to synaptic pruning, microglial phagocytosis is critical to maintaining CNS homeostasis. Consequently, it is not surprising that microglia possess several surface receptors that can trigger phagocytic activity, including TAM receptors (*45*), scavenger receptors (*46*), complement receptors (*47*), P2Y6 (*48*), and TREM2 (*11*). In regards to TREM2 specifically, it has been linked to several phagocytosis related pathways including PI3K/AKT (*49*). Moreover, TREM2 is a suggested receptor for phosphatidylserine (PS), an “eat-me” signal presented by apoptotic cells, including neurons damaged by seizure activity (*11, 50*). The elimination of neurons by microglia following seizures may even extend to neurons that are only temporarily displaying PS (*50*). With regards to our results, we observed reduced engulfment of PSVue^+^ cells when TREM2 is deficient. NeuN staining also indicated reduced neuronal engulfment by TREM2 KO microglia compared to controls, which is consistent with reduced CD68 expression. Consistently, phagocytic capacity of TREM2 KO microglia was impaired *in vitro.* Therefore, our results clearly demonstrate that phagocytic activity following IA-KA is hampered by TREM2 KO, specifically at later time points. This is in line with other studies that show TREM2 deficiency suppresses the phagocytic ability of microglia in experimental stroke (*51*). As with our other results, differences in phagocytic capacity only become significant at later timepoints, despite there being no apparent effect on the initial recognition of damaged neurons by microglia. However, like with proliferative signaling, other phagocytosis-related receptors such as TAM receptor MERTK and P2Y6, may be more influential early on (*52, 53*). This notion is supported by our RNAseq results showing that, of the mentioned genes, only TREM2 gene expression was up-regulated over baseline at 7 days after KA-induced seizures. As such, TREM2 may not be critical to the initiation of phagocytic processes but is required to complete them. Nevertheless, this effect upon the proper clearance of damaged neurons may also contribute the differences we observed in spontaneous recurrent seizure generation and frequency.

### Reduced phagocytosis may exacerbate the presentation of SRS in mice and FBTC seizures in humans

Our results suggest that the observed increase in SRS development within TREM2 KO mice could result from a combination of factors. For instance, the differences between TREM2 KO and control epileptiform patterns observed during SE seem to extend into the chronic period. What is curious, is that this maintained difference is primarily due to TREM2 KO EEG patterns, as controls returned relatively close to their genetic baseline. While not necessarily the only means by which TREM2 KO can affect the development of SRS, our results suggest that the improper clearance of dead and dying neurons from the hippocampus is a significant contributor. This may lead to neurotoxic and inflammatory cellular debris remaining in the microenvironment. Inflammation is a well-known inducer of stronger and more frequent seizures (*54–56*). Another possibility is that damaged, but not apoptotic, neurons remain within neuronal circuits inducing detrimental effects. Our results with human tissues may support the possibility that improper severance of damaged neuronal circuits increases the likelihood of generalized seizures. While CD68 expression was predominantly observed in temporal lobe white matter, it is possible that in humans, circuit regulation is conducted via severance of nerve fiber connections rather than complete phagocytosis of neurons like in mice. However, the result would be similar, higher phagocytic activity reduces the chance for focal dysfunction to develop into generalized seizures. Of the studies that explored the ability of microglia to influence SRS and chronic epilepsy, most focused on modulation of inflammation (*57*). Unsurprisingly, most studies found that inhibition of microglia inflammatory effects was beneficial. However, we recently demonstrated that in addition to being detrimental within SE, microglial depletion was also detrimental to the development of SRS (*5*). Thus, microglia may have protective roles within chronic seizures and epilepsy.

Our results clearly demonstrate that reduced microglial phagocytic activity broadly, and TREM2 KO specifically, exacerbates generalized FBTC seizure development. Thus, it is possible that enhanced stimulation of TREM2 signaling may be beneficial to seizure recovery and epilepsy. Recently, the agonistic TREM2 antibody AL002a was shown to have therapeutic potential within both multiple sclerosis (MS) and AD (*58, 59*). Additionally, other agonistic TREM2 antibodies have been developed that can specifically target human TREM2 (*60*). These antibodies, or similar therapeutics, could be used as seizure and epilepsy interventions, if enhancement of phagocytic activity is indeed advantageous. This is particularly important given the known association of drug resistant TLE and prior prolonged febrile convulsions in infancy. It has long been thought that a single prolonged febrile convulsion can initiate hippocampal epileptogenesis that years later manifests as drug resistant TLE (*61, 62*). However, like with other aspects of microglial activity, whether or not microglial phagocytic response is beneficial or detrimental to epilepsy pathogenesis is still up for debate, with it likely being context, stage, and brain region dependent. Consequently, further investigation is needed to tease out the effect microglia have upon seizures and epilepsy, both beneficial and detrimental. Regardless, our study indicates that microglial TREM2 offers a possible target for therapeutic intervention.

## MATERIALS AND METHODS

### Study Design

The overall objective of this study was to determine the role microglial TREM2 played within seizure and epilepsy development. We determined that TREM2 deficiency resulted in lower microglial phagocytic activity and enhanced seizure frequency and development. We also investigated the link between phagocytic activity and generalized seizure development within humans. For animal experiments, age matched male mice were used for each investigation. Sample exclusion was performed using the following criteria, animals needed to present 5+ FBTC events during the initial SE period and survive for 7+ days post-KA to be enrolled in subsequent experiments. A description of sample size (N) is provided in the figure legends. For experiments utilizing cell culture, 2 independent experiments were performed. For experiments utilizing human tissues all samples were obtained from the Mayo Clinic Tissues Registry. Investigators were not blinded during data collection or analysis.

### Animals

6-10 week old male TREM2^-/-^:CX3CR1^GFP/+^ (TREM2 KO) and CX3CR1^GFP/+^ (Control) mice were used for all experiments in accordance with the guidelines set forth by the Mayo Clinic Institutional Animal Care and Use Committee. Parental TREM2^-/-^ mice were obtained from Dr. Marco Colonna at Washington University School of Medicine in St. Louis, Missouri. To generate the functional general KO, portions of the transmembrane and cytoplasmic domains were deleted (*63*). Parental CX3CR1^GFP/GFP^ (B6.129P2(Cg)-Cx3cr1^tm1Litt^/J; 005582) and wild-type (C57BL/6J; 000664) mice were obtained from Jackson Labs (Bar Harbor, ME). All mice were housed in a temperature-controlled facility on a 12h day/night cycle and provided food and water *ad libitum*.

### Seizure Induction

To induce seizures, we used the intra-amygdala (IA) kainic acid (KA) administration model (*18*). Briefly, a guide cannula (-0.94 AP, -2.85 ML) was implanted 7 days before administration of 0.2 μL KA (0222; Tocris, Minneapolis, MN; 1.5 mg/mL; -3.75 DV) in PBS. Seizure status was terminated 45 mins after the initial FBTC event using valproic acid (P4543; MilliporeSigma, Burlington, MA; 300 mg/kg; *i.p.*). Seizure behaviors were monitored using a modified Racine scale (*43*). A score of 4+ was recorded as a FBTC event.

### Electroencephalogram (EEG) Evaluation

For animals undergoing EEG evaluations a single channel cortical electrode was implanted concurrently with the IA guide cannula. EEG evaluations were conducted using EEG100C amplifiers and MP150 digitizing equipment (BioPac, Goleta, CA) and synchronized with parallel video monitoring as previously described (*64*). Animals that met inclusion criteria then underwent continuous 24/7 video EEG monitoring to assess spontaneous recurrent seizure (SRS) development starting at day 7 post-KA. Baseline recordings were made 7 days post electrode and cannula implantation on animals that did not receive KA administration.

### EEG Analysis

EEG data was analyzed using custom algorithms (Matlab version 2019b) for characterizing the changes of EEG power in spectral bands of interest (Delta (1 -3 Hz), Theta (3 – 9 Hz), Alpha (8 - 12 Hz), Sigma (12-15 Hz), Beta (15 – 30 Hz), Low Gamma (30 -55 Hz), High Gamma (65-115Hz)) as described previously (*65*). Absolute and relative power spectral densities (PSD) across frequency bands were calculated using 10 second windows (epochs) and the average of PSD from all epochs was calculated as a representative value of PSD for each phase of experiment for each subject.

### Tissue Collection

Tissue was collected as previously described (*13*). 16 μm tissues sections were prepared using a cryostat (Leica Microsystems, Deerfield, IL). Sections were stored at -20°C until use.

### BrdU Administration and Tissue Preparation

The thymidine analog bromodeoxyuridine (BrdU) (B5002; MilliporeSigma) was used to label proliferating and recently post-mitotic cells as previously described (*66*). BrdU solution was intraperitoneally (*i.p*.; 100 mg/kg) injected twice per day for different periods after KA injection.

### Immunofluorescent Staining

Immunofluorescence staining was conducted as previously described (*13*). Staining was conducted overnight at 4°C with anti-NeuN (1:500; ab104225; Abcam, Waltham, MA), rabbit anti-CD68 (1:500; ab125212; Abcam), rat anti-BrdU (1:500; ab6326; Abcam) and/or rabbit anti-Iba1 (1:500; ab178847; Abcam) primary antibodies. Sections were then incubated with appropriate secondary antibodies for 60 mins at room temperature (1:500; Alexa Fluor goat anti-rabbit 594, A11037; Alexa Fluor goat anti-rabbit 488, A11008; Alexa Fluor goat anti-rat 594, A11007; Invitrogen, Waltham, MA) before mounting with DAPI Fluoromount-G (0100-20; SouthernBiotech, Birmingham, AL). Fluorescent images were captured with either an EVOS fluorescent microscope (ThermoFisher Scientific, Waltham, MA) or BZ- X800E fluorescent microscope (Keyence, Itasca, IL). Image analysis was performed using FIJI (ImageJ/Fiji, version 1.53o; National Institutes of Health, Bethesda, MD).

### *In Vitro* Phagocytosis Assay

Briefly, primary mixed glial cultures were isolated from P2-P5 control and TREM2 KO mice as previously described (*67*). Cultures were maintained at 37°C, 5% CO_2_ in DMEM:F12 containing 1% pen-strep and 10% FBS on 5 poly-D-lysine (PDL; P7405- 5MG; Sigma-Aldrich, St. Louis, MO) coated T-75 flasks. Cultures were then maintained for 10- 12 days, with media changes conducted every 3-4 days. Primary microglia were then isolated by shaking flasks on an orbital shaker for 30 mins at 200 RPM and plated into PDL coated 24-well plates at a density of 50K per well. Isolated microglia were maintained in astrocyte conditioned DMEM:F12 to maintain viability.

To generate pHrodo Red labeled apoptotic N2a cells, N2a cultures were first incubated for 16-20 h in EMEM containing 1% pen-strep and 10% FBS supplemented with 0.5 µM staurosporine (ab146588; Abcam) then labeled with the pHrodo Red Cell Labeling Kit for Incucyte (4649; Sartorius, Bohemia, NY) with a final concentration of 125 ng/mL dye following manufacturer’s instructions. Labeled apoptotic N2a cells were then added to the primary microglia cultures at a ratio of 1:1. Well plates then immediately underwent imaging with an Incucyte SX5 equipped with a SX5 G/O/NIR optical module (Sartorius). Quantifications were conducted with the systems onboard analysis software.

### PSVue Administration

*In vivo* PSVue administration was conducted in a similar manner as previously described (*35*). Briefly, 3 µL of 1mM PSVue 550 solution (P-1005; Molecular Targeting Technologies, West Chester, PA) was injected via an implanted intracranioventricular (ICV) cannula 3 h prior to sacrifice.

### Analysis of Microglial Engulfment of PSVue Positive Cells

Brain slices were imaged with a Zeiss LSM980 for confocal imaging (Carl Zeiss, Oberkochen, Germany) through a 63× oil-immersion objective lens. GFP^+^ cells and PSVue^+^ cells were characterized from 3D image data (1024×1024 pixels, 0.044 μm/pixel, 0.24 μm Z-step, 50 slices). To measure the fluorescent intensity of the GFP signal, an outline region of interest (ROI) was manually created using “free hand selection” tool in FIJI (ImageJ/Fiji, version 1.53o; NIH). The ROIs were created around an average of 30 PSVue^+^ cells per mouse. The image plane with the largest cell diameter was used to draw the ROI for each PSVue^+^ cell. Coverage was calculated as a percentage of the ROI containing GFP signal. GFP positive contact was defined by pixels with a signal intensity value greater than the mean + 2SD of ROIs around PSVue^+^ cells without GFP^+^ cell contact (baseline) in the same brain slice.

### Fluoro-Jade C (FJC) Staining

Sections were stained with FJC (AG325, Millipore-Sigma, Burlington, MA) in 0.1% acetic acid for 30 mins following manufacturer’s instructions. Briefly, sections were washed twice with ddH_2_O, 1% NaOH in 80% EtOH for 5 mins, 70% EtOH for 2 mins, and then ddH_2_O for 2 min. Sections where then incubated in freshly prepared 0.06% potassium permanganate solution 30 mins, washed with ddH_2_O, incubated with FJC (0.0001% working solution) for 30 mins protected from light, then washed again with ddH_2_O. Sections were then cleared with xylene and mounted with DPX mounting media (VWR, Hatfield, PA). Fluorescent images were captured with an EVOS fluorescent microscope (ThermoFisher Scientific, Waltham, MA). Image analysis was performed using FIJI (ImageJ/Fiji, version 1.53o; National Institutes of Health, Bethesda, MD).

### RNA Extraction

Immediately following transcardiac perfusion with 50 mL PBS, brains were removed and the ipsilateral area around the previously defined coordinates for KA injection was isolated. Tissues collected from 2-4 mice were pooled per sample. Samples then underwent cellular isolation via CD11b magnetic bead sorting using the MACS Adult Brain Dissociation kit (130- 107-677; Miltenyi Biotec, Gaithersburg, MD) per manufacturer’s instructions. Total RNA was then extracted using the RNeasy Microkit (74004; Qiagen, Germantown, MD). RNA purity and integrity was assessed via Eukaryote Total RNA Pico Chip (Agilent Technologies, Santa Clara, CA). RNA was stored at -80°C until use.

### RNAseq Library Preparation and Analysis

RNAseq library preparation and initial analysis was performed by BGI (Cambridge, MA) using the DNBSEQ platform (MGI Tech Co., Ltd., Shenzhen, China). Raw sequencing data filtering, alignment (aligned to the mus musculus GCF_000001635.27_GRCm39 reference genome), and expression determination was also conducted by BGI. DESeq2 was used to determine differentially expressed genes (DEGs) between conditions (*68*). A Q value (adjusted P value) ≤ 0.05 was used to determine significance between conditions. Additional analysis was performed using the Dr. Tom data visualization platform (BGI).

### Human Tissue Analysis

Formalin fixed paraffin embedded (FFPE) human tissue sections were obtained from the Mayo Clinic Tissues Registry and all related experiments were approved by the Mayo Clinic Institutional Review Board (IRB; ID:21-012340). Material was obtained from patients with mesial temporal lobe focal intractable epilepsy who elected to undergo neurosurgical administration (Patient N = 31). Included cases had no known history of focal cortical dysplasia, primary CNS tumor, or metastatic brain disease. Both male (41.9%) and female (58.1%) patients ranging in age 6 to 70 at the time of resection (average age 38.0) were utilized. Patients were evaluated by the specialized epilepsy clinic in the Neurology Department of the Mayo Clinic and all patients underwent comprehensive evaluation including MRI and EEG studies. Briefly, tissue sections were first deparaffinized and rehydrated prior to incubation with 0.01% Sudan Black for 7 mins. Antigen retrieval was performed via citrate buffer treatment (H-3300-250; Vector Laboratories, Newark, CA) and then sections underwent immunofluorescent staining as outlined previously. Sections were incubated overnight at 4°C with rabbit anti-Iba1 (1:500; ab178847; Abcam) and mouse anti-CD68 (1:200; M0814, clone KP1; Agilent Technologies, Santa Clara, CA) primary antibodies. Sections were then incubated with secondary antibodies (1:500; Alexa Fluor goat anti-rabbit 488, A11008; Alexa Fluor goat anti-mouse 594, A11032; Invitrogen), and mounted with DAPI Fluoromount-G. Fluorescent images were captured with an EVOS fluorescent microscope. Image analysis was performed using FIJI (ImageJ/Fiji, version 1.53o; NIH).

### Statistical Analysis

All results are displayed as Mean ± SEM. Statistical analysis was performed with GraphPad Prism software (version 9; San Diego, CA). Comparisons between two groups were performed using Unpaired T-test with Welch correction unless otherwise stated. P values < 0.05 were considered as statistically significant.

## Supporting information

All supplements

## Acknowledgments

The authors would like to thank Drs. Nathan Staff and Sybil Hrstka (Mayo Clinic) for the assistance with the Sartorius Incucyte system. We thank Dr. Vanda Lennon (Mayo Clinic) for critical reading of the manuscript.

National Institute of Health grant R35NS132326 (LJW)

National Institute of Health grant R01NS088627 (LJW)

National Institute of Health grant R01NS112144 (GAW, LJW)

## Author Contributions

Conceptualization: DBB, LJW

Methodology: DBB, VK, KH, ANT, GAW, LJW

Investigation: DBB, VK, KH, SZ, LW, BAE, AD, JZ, JFP, MX, LW

Visualization: DBB, VK, KH, LW

Funding acquisition: LJW

Supervision: LJW

Writing – original draft: DBB, LJW

Writing – review & editing: DBB, BAE, ANT, GAW, LJW

## Data and materials availability

All data will be maintained within the corresponding author’s laboratory and will be freely available upon request.

