## Supplementary material for "Impaired microglial phagocytosis promotes seizure development": All supplements

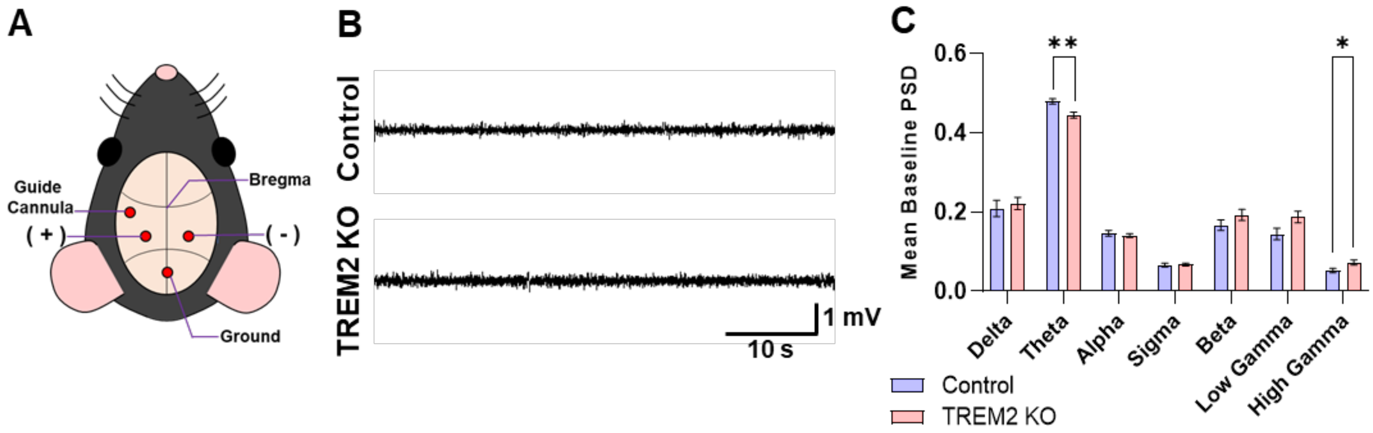


**Fig. S1: Additional EEG analysis comparing TREM2 KO and control. (A)** Schematic of IA guide canula and EEG electrode placement. **(B)** Representative baseline EEG profiles of TREM2 KO and control mice. EEG profiles for each animal were investigated for differences in individual frequency band power spectral density (PSD). Delta: 1-3 Hz; Theta: 3-9 Hz; Alpha: 8-12 Hz; Sigma: 12-15 Hz; Beta: 15-30 Hz; Low Gamma: 30-55 Hz; and High Gamma: 65-110 Hz. **(C)** Baseline PSD values. Data presented as Mean ± S.E.M. $N \leq 10$ for each group. Unpaired T test with Welch correction was used for each comparison. * $P < 0.05$, ** $P < 0.01$, *** $P < 0.001$.

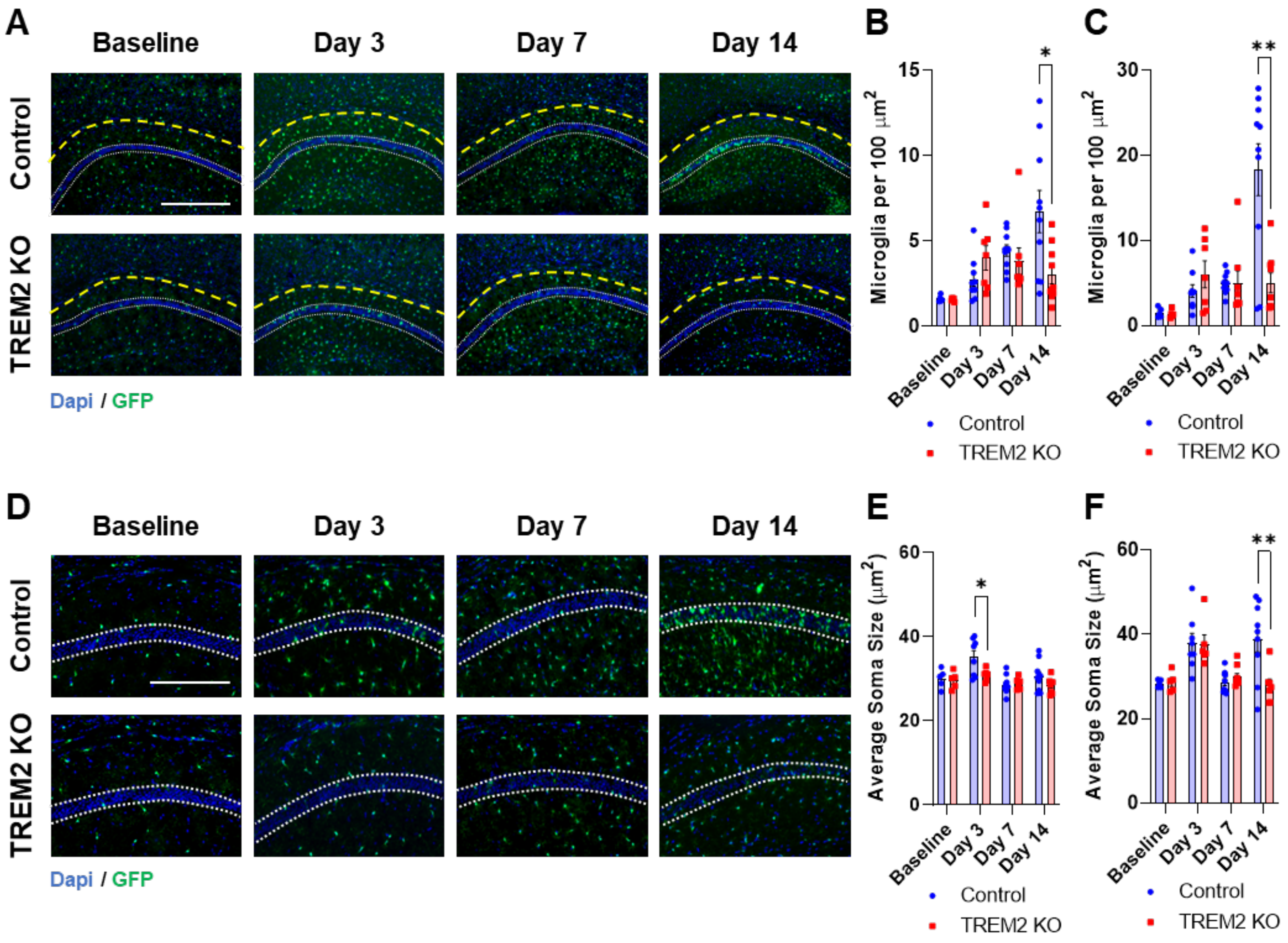


**Supplemental Fig 2: Effect of TREM2 KO on microglial number within the CA1 region following IA-KA. (A)** Representative images of the CA1 region at baseline and 3, 7, 14 days post-KA administration. Scale bar = 400 µM. **(B)** Quantification of microglia density within the imaged CA1 region (below the yellow dashed line). **(C)** Quantification of microglia density within the *stratum pyramidale* (between white dashed lines). **(D)** Representative images of the CA1 region at baseline and 3, 7, and 14 days post-KA administration. Scale bar = 200 µm. **(E)** Quantification of microglial soma size within the image field. **(F)** Quantification of microglial soma size within the *stratum pyramidale* (between white dashed lines). Data presented as Mean ± S.E.M. N = 5-10 for each group. Image fields collected from 2-3 non-consecutive tissue sections were averaged for each N. Unpaired T test with Welch correction was used for each comparison at each time point. * $P < 0.05$, ** $P < 0.01$

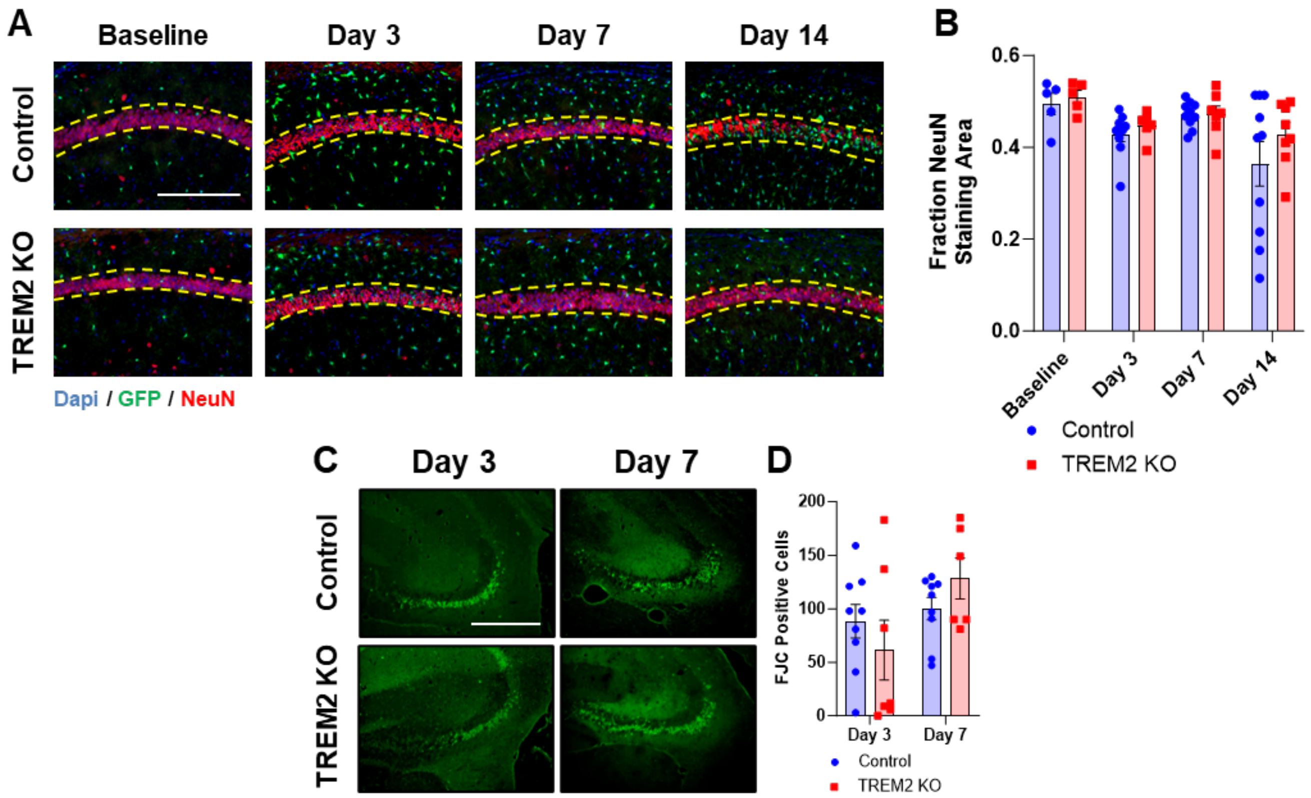


**Fig. S3: Neuronal atrophy following IA-KA administration. (A)** Representative images of NeuN staining within the CA1 region at baseline and 3, 7, 14 days post-KA administration. Scale bar = 200 µm. **(B)** Quantification of NeuN staining. Stained area divided by total area (between yellow dashed lines). (**C**) Representative images of Fluoro-Jade C (FJC) positive neurons within the CA3 region. (**D**) Quantification of FJC positive neurons. Data presented as Mean ± S.E.M. N = 5-10 for each group. Image fields collected from 2-3 non-consecutive tissue sections were averaged for each N. Unpaired T test with Welch correction was used for each comparison at each time point. * $P < 0.05$, ** $P < 0.01$

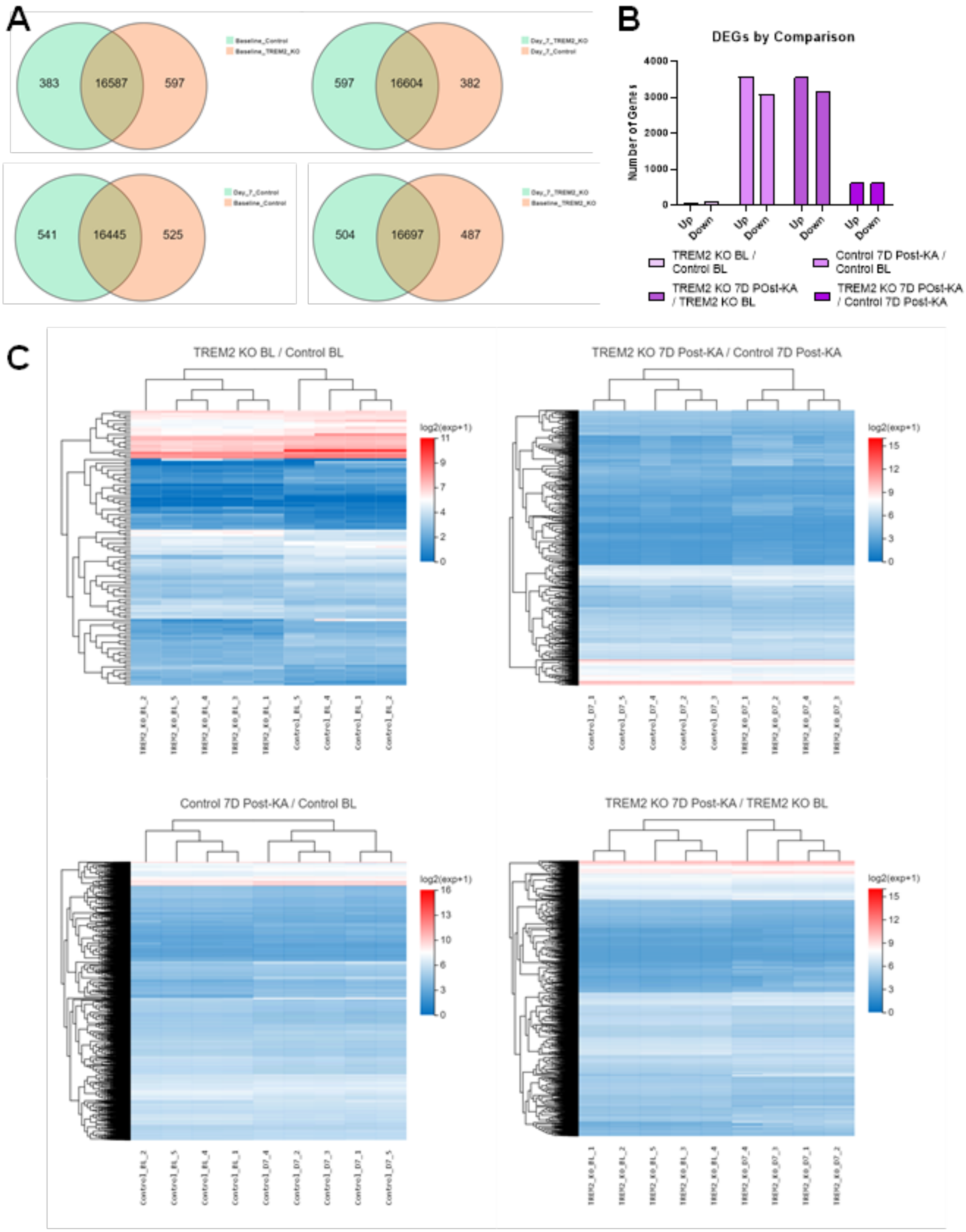
A
Baseline_Control
Baseline_TREM2_KO
383
16587
597
Day_7_TREM2_KO
Day_7_Control
597
16604
382
Day_7_Control
Baseline_Control
541
16445
525
Day_7_TREM2_KO
Baseline_TREM2_KO
504
16697
487
B
DEGs by Comparison
Number of Genes
4000
3000
2000
1000
0
Up
Down
Up
Down
Up
Down
Up
Down
TREM2 KO BL / Control BL
Control 7D Post-KA / Control BL
TREM2 KO 7D POst-KA / TREM2 KO BL
TREM2 KO 7D POst-KA / Control 7D Post-KA
C
TREM2 KO BL / Control BL
log2(exp+1)
11
9
7
4
2
0
TREM2_KO_BL_2
TREM2_KO_BL_5
TREM2_KO_BL_4
TREM2_KO_BL_3
TREM2_KO_BL_1
Control_BL_5
Control_BL_4
Control_BL_3
Control_BL_2
TREM2 KO 7D Post-KA / Control 7D Post-KA
log2(exp+1)
15
12
9
6
3
0
Control_D7_1
Control_D7_5
Control_D7_4
Control_D7_2
Control_D7_3
TREM2_KO_D7_1
TREM2_KO_D7_2
TREM2_KO_D7_4
TREM2_KO_D7_3
Control 7D Post-KA / Control BL
log2(exp+1)
16
13
10
6
3
0
Control_BL_2
Control_BL_5
Control_BL_4
Control_BL_1
Control_D7_4
Control_D7_2
Control_D7_3
Control_D7_1
Control_D7_5
TREM2 KO 7D Post-KA / TREM2 KO BL
log2(exp+1)
15
12
9
6
3
0
TREM2_KO_BL_1
TREM2_KO_BL_2
TREM2_KO_BL_5
TREM2_KO_BL_3
TREM2_KO_BL_4
TREM2_KO_D7_4
TREM2_KO_D7_3
TREM2_KO_D7_1
TREM2_KO_D7_2

**Fig. S4: Overview of expression pattern differences between TREM2 KO and control. (A)** Venn diagrams illustrating the overlap between identified gene lists. **(B)** Number of differentially expressed genes (DEGs) by comparison. **(C)** Heatmaps illustrating the relative expression profiles for each comparison.

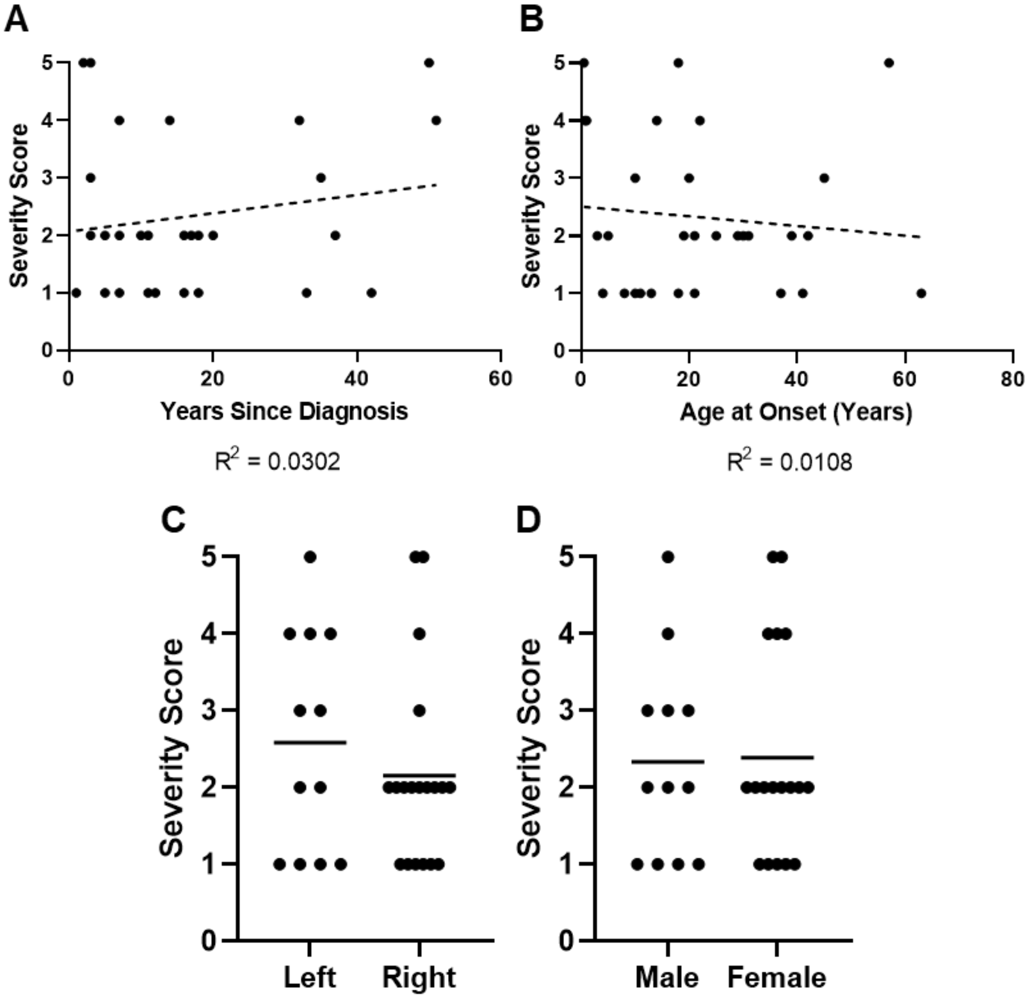


**Fig. S5: Relationship between severity score and patient demographic information.** No relationship was observed between severity score and **(A)** the amount of time since diagnosis or **(B)** the patient age at onset. There was also no relationship between **(C)** which brain hemisphere resections were taken, or **(D)** the sex of the patient.
